# Transcription factor TFII-I fine tunes innate properties of B lymphocytes

**DOI:** 10.1101/2022.09.21.508949

**Authors:** Amit Singh, Mary Kaileh, Supriyo De, Krystyna Mazan-Mamczarz, Dashzeveg Bayarsaihan, Ranjan Sen, Ananda L Roy

## Abstract

The ubiquitously expressed transcription factor TFII-I is a multifunctional protein with pleiotropic roles in gene regulation. TFII-I associated polymorphisms are implicated in Sjögren’s syndrome and Lupus in humans and, germline deletion of the *Gtf2i* gene in mice leads to embryonic lethality. Here we report a unique role for TFII-I in homeostasis of innate properties of B lymphocytes. Loss of *Gtf2i* in murine B lineage cells leads to a change in transcriptome and chromatin landscape, which resembles myeloid-like features and coincides with enhanced sensitivity to LPS induced transcription. TFII-I deficient B cells also show increased switching to IgG3, a phenotype associated with inflammation. These results demonstrate a role for TFII-I in maintaining immune homeostasis and provide clues for *GTF2I* polymorphisms associated with B cell dominated autoimmune diseases in humans.

## Introduction

TFII-I was discovered as a transcription factor that bound to the adenovirus major late core promoter Initiator (Inr) element. TFII-I also interacts with a sequence-specific DNA element (the E-box element) together with the upstream stimulatory factor (USF) in cell-free systems (*1*). Subsequent studies have suggested a broader, multifunctional role for TFII-I in a variety of cell types perhaps by virtue of its interaction with cell type-specific proteins and signaling intermediates (*2*). TFII-I is shown to be phosphorylated in response to several cell surface receptor signaling pathways, including growth factor receptor and immune receptors (*3*), further suggesting its role in these pathways. Recent studies also indicate its involvement in DNA-damage and response pathways (*4*).

TFII-I is vertebrate-specific and encoded by the *GTF2I* gene in humans and *Gtf2i* in mice (*5*). The locus comprises 36 exons that give rise to several alternatively spliced isoforms (*6*). Various disease pathologies are associated with either gene dosage effects or mutations or gene fusion events involving *GTF2I* (*7*). Most notably, *GTF2I* haploinsufficiency is associated with Williams-Beuren Syndrome (WBS) a neurodevelopmental disorder with specific craniofacial features (*7-9*). *GTF2I* gene dosage is further ascribed to autism spectrum disorder (ASD) (*10*) and single nucleotide polymorphisms in *GTF2I* loci are associated with autoimmune disorders like SLE, RA and Sjogren syndrome, reviewed in (*3*). Finally, a single point mutation in *GTF2I* is associated with thymic epithelial tumors as well as *GTF2I* gene fusions are known to occur in various forms of cancers (*11, 12*). Collectively, these studies clearly point to an essential and important role for TFII-I in various human diseases. Consistent with pleiotropic roles of TFII-I, germline deletion of the gene *Gtf2i* in mice leads to early embryonic lethality likely due to severe defects in vasculogenesis and angiogenesis (*13*).

Although ubiquitously expressed in most if not all cell types, a series of studies showed that TFII-I biochemically interacts with Bruton’s tyrosine kinase (Btk) in B cells and corresponding kinase Itk in T cells, suggesting a role in immune cell type specific functions (*14*). Moreover, biochemically TFII-I interacts with the B-cell specific co-activator, OCA-B (*15*) and B cell transcription factor Bright (*16*) and regulates Immunoglobulin gene expression in cell-based assays (*15*). Intrigued by TFII-I’s potential involvement in the immune system, we conditionally deleted the *Gtf2i* gene in mice using a CD19-Cre driver to test its potential role in mature B cells.

Murine naïve B cells are generally classified into three distinct subsets, B-1 B cells of peritoneal origin, follicular (FO) B cells, and marginal zone (MZ) B cells (*17*). A large fraction of FO B cells is IgD^hi^IgM^low^CD21^mid^ cells (termed as follicular type I B cells), while a smaller fraction are IgD^hi^IgM^hi^CD21^mid^ B cells (termed as follicular type II B cells (*18*). In contrast, only a minor population of splenic B cells are marginal zone (MZ) cells expressing high levels of IgM, CD21 and CD1d. MZ B cells are generated as naive B cells that intrinsically have some properties resembling those of memory cells. MZ B cells are also considered to be innate-like cells that can be induced to differentiate into short-lived plasma cells in the absence of BCR ligation (*19*). These properties allow the MZ B cells to crossover between adaptive and innate immunity (*20*).

We show here that ablation of TFII-I in B cell lineage selectively reduces murine MZ B cells. Further, transcriptomic and chromatin studies show that the surviving MZ B cells as well as FO B cells exhibit enhanced innate cell-like molecular features. Consistent with this notion, the splenic B cells in TFII-I deleted mice exhibit a noticeable increase in lipopolysaccharide (LPS) sensitivity, once again resembling features that are characteristic of innate immune cells. These results demonstrate that TFII-I is critical for maintaining B cell homeostasis and its absence accentuates innate properties of B lymphocytes.

## Results

### Reduced numbers of MZ B cells in *Gtf2i*^*fl/fl*^*CD19-Cre*^*+*^ (*Gtf2i* cKO) mice

To interrogate the role of TFII-I in B-cell function we generated a “conditional knock-out” *Gtf2i* mouse model by breeding ‘floxed’ alleles of *Gtf2i* (*21*) with B cell-specific CD19-driver Cre. The B cell specific deletion of *Gtf2i* at genomic, mRNA and protein level were confirmed (Supp Fig. 1A-B). Total splenocytes and B cell numbers were unaffected by *Gtf2i* ablation (Supp Fig. 1C). Various splenic B cells subsets (transitional, FO and MZ) were identified by previously described flow cytometry gating strategy (*17*) (Fig. 1A) and the percentages of total B, FO, MZ and transitional B (T1, T2) cells from 18 - 20 independent experiments are shown (Fig. 1). There was no appreciable difference in the percentages of splenic B cells and T1, T2 cells between control (wild type, WT and heterozygous, Het) and *Gtf2i* cKO (homozygous, HO) mice (Fig. 1B and 1C). FO B cells percentage was slightly increased in cKO compared to control groups (Fig. 1D). However, MZ B cells proportions were significantly reduced in cKO compared to WT control mice (Fig. 1E).

**Figure 1:**
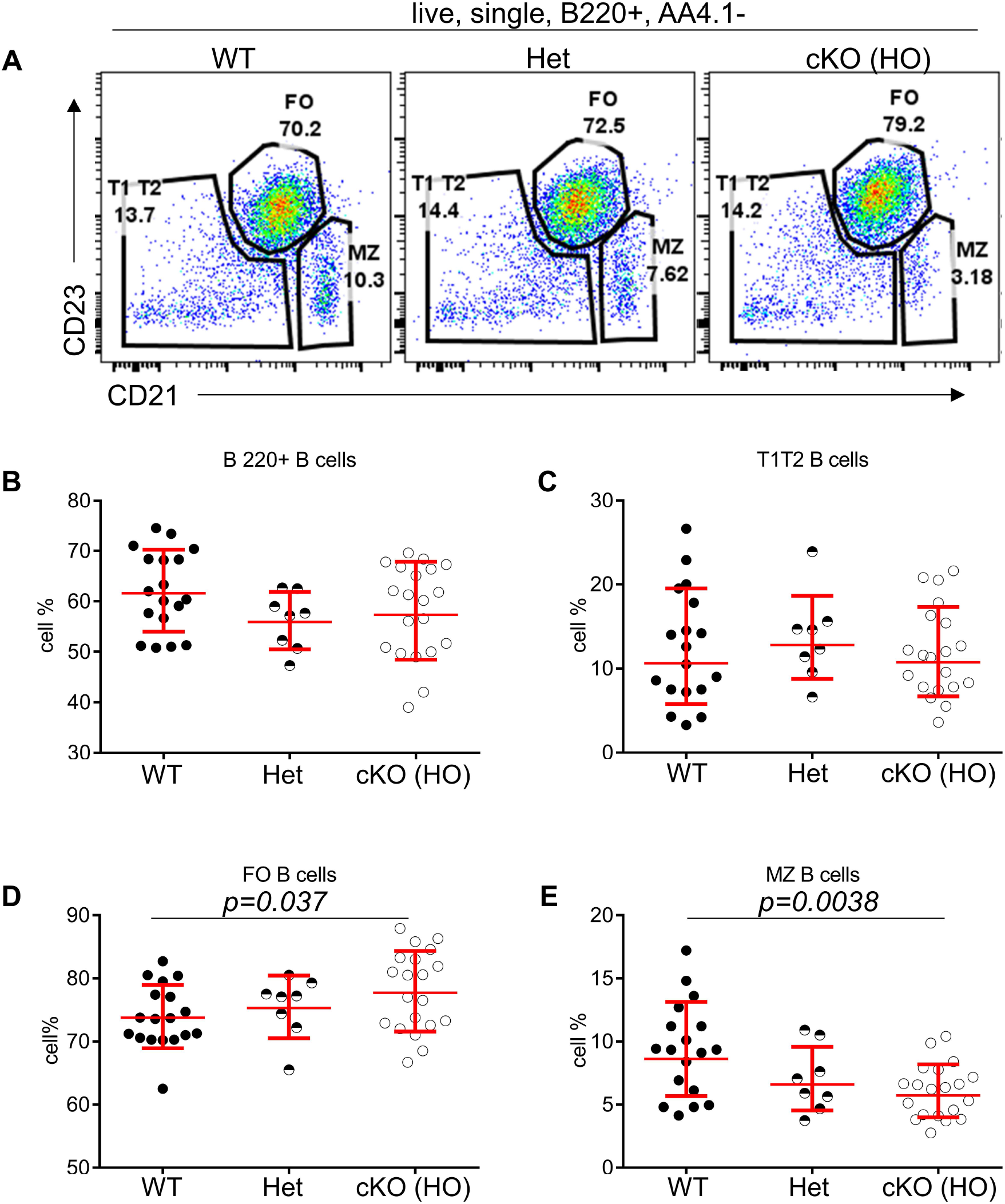
Reduced percentages of MZ B cells in *Gtf2i* cKO mice: (A) Representative plots of splenic B cell immunophenotyping of WT, *Gtf2i*^*fl/+*^*CD19-Cre+* (Het) and *Gtf2i*^*fl/fl*^*CD19-Cre+* (HO) mice. FACS dot plots of CD21 (x axis) and CD23 (y axis) expression in B cells show CD21^lo^CD23^hi^ Follicular (FO) B cells, CD21^hi^CD23^lo^ Marginal Zone (MZ) B cells and CD21^lo^CD23^lo^ Transitional (T1T2) B cells. (B) Splenocytes of wild type (WT), heterozygous (Het) and homozygous null (HO) mice were stained with B220 antibody and the percentages of B220+ total B cells are shown. (C – E) Primary splenic B cells were isolated from WT, Het and HO mice using EasySep magnetic purification techniques by negative selection. About 1 million B cells were stained with anti CD21-FITC and anti CD23-PE for 35 minutes, then washed and analyzed at the BD FACSAria Fusion. The percentages of FO, MZ and T1T2 B cells from 18 - 20 independent experiments are shown. P values were calculated using paired student t test in prism.

We further investigated the tonic signaling (low level of antigen independent BCR signaling responsible for B cell survival) (*22*) in total splenic B cells in absences of *Gtf2i*, as it can have Btk mediated B cell survival effects (*23, 24*). B cells were cultured *ex vivo* with or without B cell activating factor (BAFF) for the indicated times (Supp Fig. 1D) and viability was checked using flow cytometry. However, there was no appreciable difference in the viability of B cells with or without BAFF among the WT, Het, and HO mice, suggesting no obvious role of TFII-I in B cell survival under these experimental conditions. Taken together these results indicate that although B cell specific ablation of *Gtf2i* does not interfere broadly with peripheral B cell homeostasis and survival, it selectively reduces MZ B cells in the spleen.

### Effects of *Gtf2i* ablation on B transcriptome

To understand the possible mechanism of peripheral differences in B cell subsets, we analyzed the transcriptome using bulk RNA-seq. Total RNA was prepared from FACS sorted FO and MZ B cells of WT and cKO mice spleens. We first identified differentially expressed genes (DEGs) from FO and MZ B cell subsets between *Gtf2i* cKO and WT mice via DESeq2 algorithm taking ≥1.5 fold and FDR≤ 0.05 as cutoff (Fig. 2 and Supp Fig. 2A). DEG numbers were low for both subsets: in FO B cell only 6 DEGs were WT specific and 22 were unique to *Gtf2i* cKO (Supp Fig. 2B), while MZ B cells showed 64 DEGs unique to WT and 40 genes specific to *Gtf2i* cKO (Fig. 2A). However, we noticed, RNA binding protein (RBP) family gene, *Zfp36l2* which is involved in MZ B cell maintenance (*25*) and regulation of immune inflammation (*26*), was selectively downregulated in cKO MZ B cells. This may partially provide an explanation as to why the cKO mice have reduced numbers of MZ B cells. Whether *Zfp36l2* is a target of TFII-I is presently unknown.

**Figure 2:**
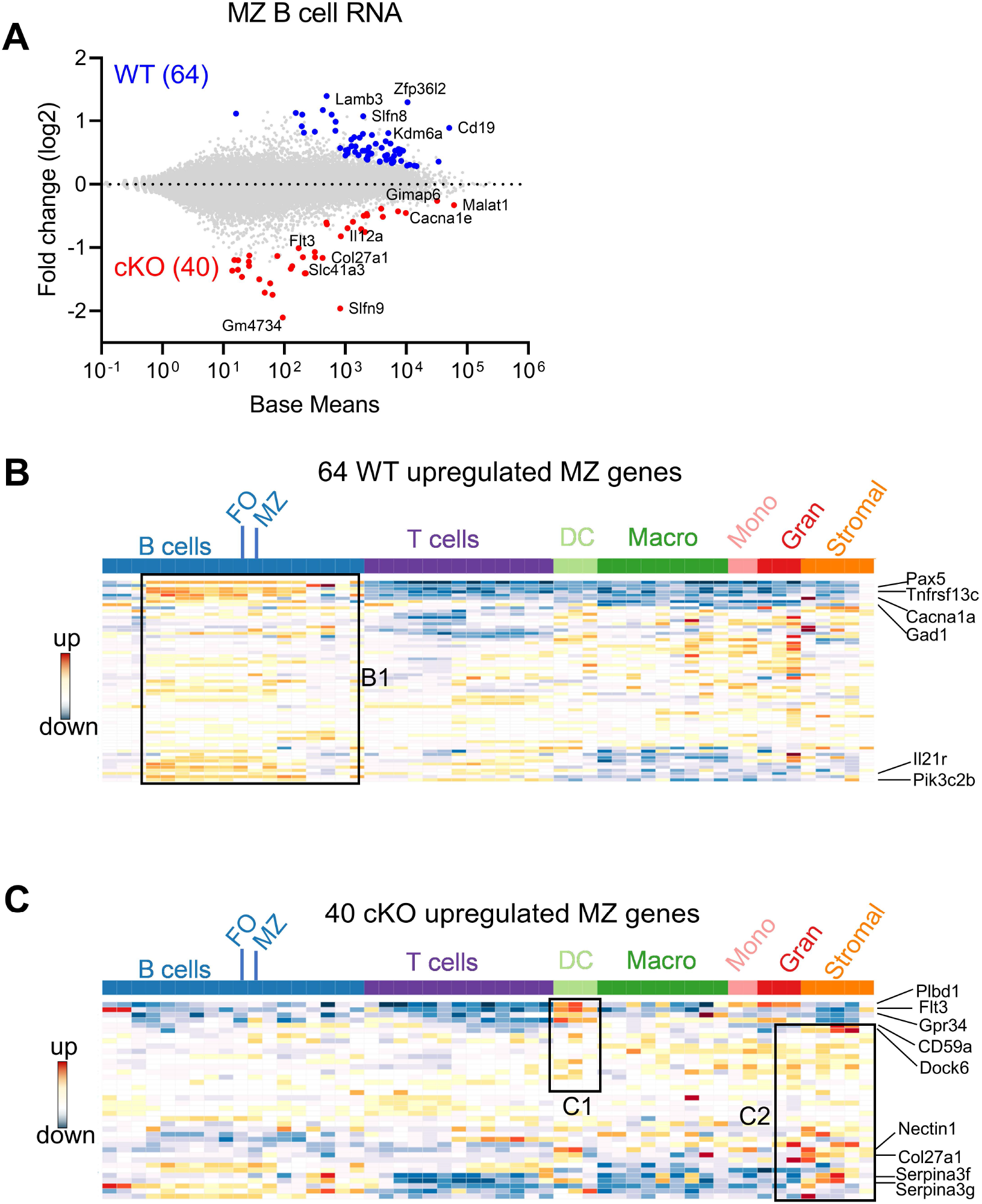
FO and MZ B cell defining transcriptome in WT and cKO (homozygous null) mice: Total RNA was extracted from FO and MZ B cells of WT and cKO mice. The RNA was used for bar-coded library preparations and sequencing. (A, B) MA plot displaying the log fold-change compared with mean expression and showing the differentially expressed genes (DEGs) in FO and MZ B cells of the WT (A) and of the cKO (B) mice. (C, F) Venn diagram showing the genes that are common verses the ones unique to FO (C) and MZ (F) B cells between the WT and the cKO mice. (D) Heatmap demonstrating average counts (CPM, n=2) normalized by DESeq2 of the 275 FO common genes (left panel) and of the 105 unique cKO genes (right panel). (E) 105 cKO FO specific genes were queried against the ImmGen-Database to identify their enrichment in the different immune cell populations. Heatmap representing the expression of these genes in all immune cells. Genes enriched in the B cell compartments are shown in box E1, genes enriched in the Dendritic and Macrophage cells are shown in box E2 while genes that are enriched in the myeloid and stromal lineages are shown in box E3 (G) Heatmap demonstrating average counts (CPM, n=2) normalized by DESeq2 of the 626 MZ common genes (right panel) and of the 201 unique cKO genes (left panel). (H) 201 cKO MZ genes were queried against the ImmGen-Database as in E and the genes that are enriched in B and T cells compartments are shown in box H1, while the genes that are enriched in Dendritic cells and Macrophages are shown in box H2 and the genes that are enriched in myeloid and stromal lineages are shown in box H3.

We next overlapped FO and MZ DEGs of cKO B cells to find common target genes of Gtf2i. Out of 62 cKO DEGs, 12 genes were shared by both FO and MZ subsets and 28 were MZ specific genes (Supp Fig. 2C). These cKO-specific genes were further showing the dominance of innate related genes (Supp Fig. 2C, boxes bold fonts). Observing these innate related genes in the absence of *Gtf2i* we further analyzed MZ DEGs for presence of potential broad immune cell features using Immunological Genome Database (ImmGen-Database) (*27*). Genes which were up-regulated with absence of TFII-I (40 genes) only demonstrate enrichment in MZ B cells (Supp Fig. 2D, box A), while genes which were downregulated in the absence of TFII-I (64 genes) were also enriched in other peripheral B cell subsets, mainly the transitional B cells genes including *Pax5, Tnfrsf13c, Cacna1, Gad1, Il21r* and *Pik3c2b* (Fig. 2B box B1 and Supp Fig. 2D, box A). Interestingly, 40 cKO genes were enriched in two splenic dendritic cell subsets of bone marrow origin (CD4^+^ and CD8^+^ DCs genes like *Plbd1, Flt3* and *Gpr34*) (*28, 29*) (Fig. 2C box C1 and Supp Fig. 2D, box B). Further, they also exhibit genes such as *Cd59a, Dock6, Nectin1, Col27a1, Serpina3f* and *Serpina3g* (Fig. 2C box C2 and Supp Fig. 2D, box C) which are transcripts from stromal subsets of Thymic Medullary Epithelial Cells “TEC”, fibroblastic reticular cells “FRC”, and lymphatic endothelial cells “LEC” origin and myeloid development genes (*30-32*). Together, these observations indicate that although *Gtf2i* deletion in B cells doesn’t make significant transcriptome alteration, it leads both FO and MZ subsets to upregulate the innate and stromal cell derived genes.

### FO and MZ B cell defining transcriptome in WT and *Gtf2i* cKO mice

Both follicular and marginal zone B cells develop from transitional B cells and attain their signature genes (*17*). Since marginal zone cells were reduced in the absence of TFII-I we hypothesized that TFII-I could alter the FO and MZ differences. To test this hypothesis, using DeSeq2 algorithm, we first identified DEGs between our WT FO-MZ B cells (Supp Fig. 3A) and compared with Immgen-Database FO-MZ DEGs. Our analysis of WT B cells identified 328 upregulated genes in FO B cells and 714 genes in MZ B cells (Fig. 3A). These numbers were similar to ImmGen-Database (*27*) FO-MZ comparison DEGs, indicating a robustness in our analysis (Supp Fig. 3B). We next assessed the FO and MZ signature DEGs in absences of *Gtf2i* using cKO mice. *Gtf2i* ablation shows slightly more DEGs between FO and MZ B cells comparable to the WT; 380 genes in the cKO FO cells, while 827 genes in the cKO MZ cells (Fig. 3B). To determine the features associated with these cKO-specific genes, they were compared to WT using Venn overlap.

**Figure 3:**
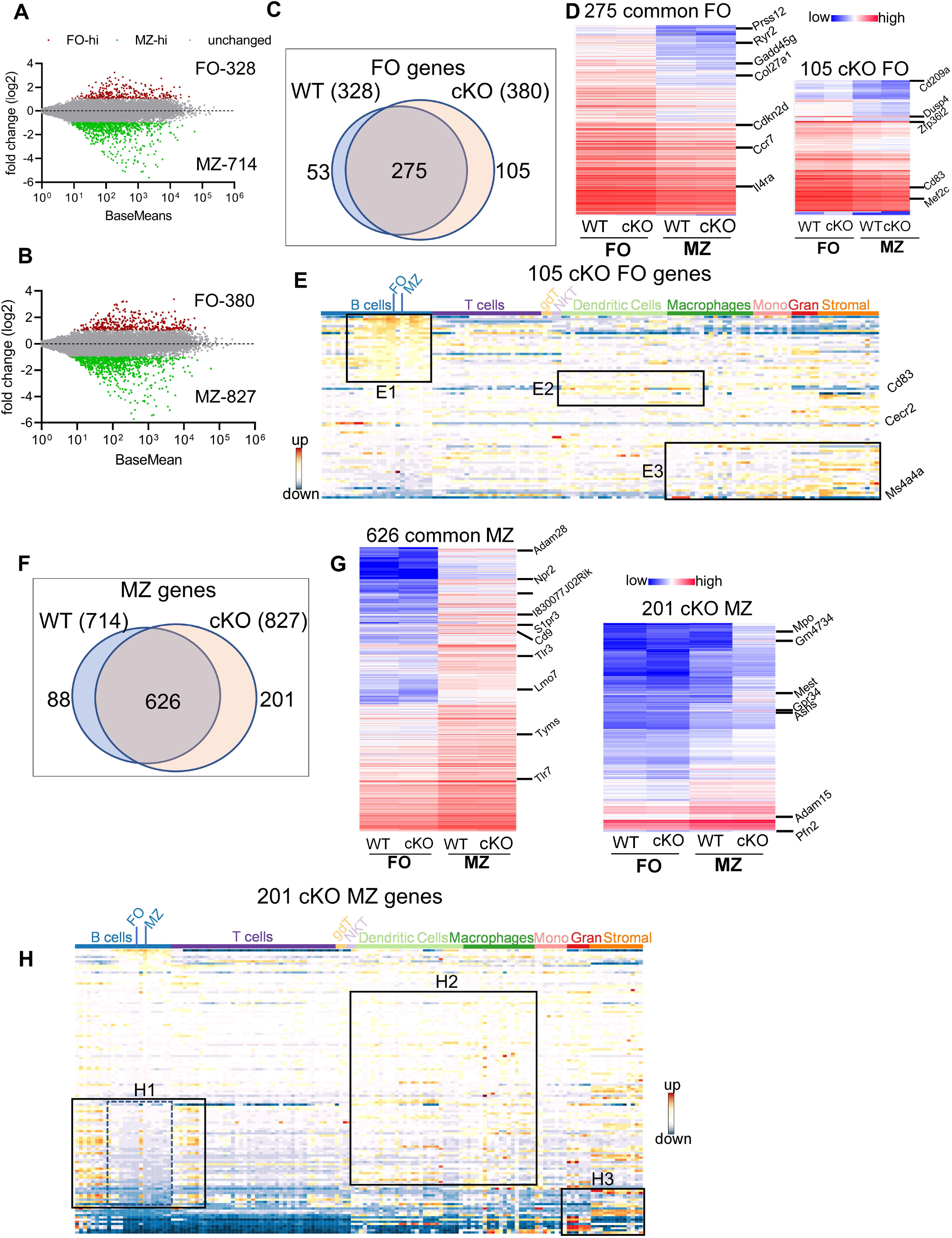
Differentially expressed genes in FO and MZ B cells derived from *Gtf2i* cKO and WT mice. Total RNA was extracted from FO and MZ B cells of WT and cKO mice. The RNA was used for bar-coded library preparations and sequencing. (A) MA plot displaying the log fold-change compared with mean expression and showing the DEGs of the WT MZ B cells (64 genes) and of the cKO MZ B cells (40 genes). (B) 64 WT MZ specific genes were queried against the ImmGen-Database and the heatmap shows the expression levels of these genes in B and other immune cells. Box B1 shows genes that are enriched in the different B cell compartments. (C) Heatmap shows the expression levels of the 40 cKO MZ specific genes when they were queried against the ImmGen-Database. Genes that are enriched in DC compartment are shown in box C1 while genes that are enriched in stromal and granulocytes are shown in box C2.

#### FO B cell genes

For FO specific genes, majority (275) were commonly upregulated (unaltered by *Gtf2i* absence) in both groups (Fig. 3C) with a similar expression level (Fig. 3D left panel and Supp Table 1). Of these genes *Prss12, Ryr2, Il4ra*, and *Gadd45g* are known to be FO specific (*33*). Top 200 common FO genes were analyzed for transcriptomic signatures of other immune cells using ImmGen-Database including B cell, T cell, dendritic cells (DC), macrophages (Macro), monocytes (Mono), granulocytes (Gran) and stromal cells (Supp Fig. 3C). These genes demonstrated features consistent with FO B cell specific genes (Supp Fig. 3C upper panel red arrow). Interestingly, nearly half of these common FO genes were also enriched in other peripheral subsets of B cells except MZ, germinal center (GC) and peritoneal cavity B cells (Supp Fig. 3D, box SD1). This suggest that transcriptionally FO B cells are significantly far from MZ, and GC B cells compare to transitional B cells. A subset of these common genes was expressed to a lower extent in mature T cells (Supp Fig. 3D, box SD2), while the genes that are expressed at lower levels in FO B cells were highly expressed in macrophages, monocytes, and stromal cells (Supp Fig. 3D, box SD3). However, 105 genes were upregulated in FO B cells in absence of *Gtf2i* including *Zfp36l2, Dusp4, Cd83, Cd209a* and *Mef2c* (Fig. 3D right panel and Supp Table 2). Similarly, 53 genes were downregulated in *Gtf2i* ablated FO B cells (upregulated in WT FO B cells) including *Dab2ip* and *Abca8b*. Immgen-Data base comparison of these 53 genes shows that they were also expressed in other B cell subsets (Supp Fig. 3E, box SE1), including *Vpreb3* (a gene expressed during B cell development) (*34*). The poorly expressed FO WT genes were expressed in other myeloid and stromal cell compartments (Supp Fig. 3E, box SE2), a feature observed in the common set of FO genes (Supp Fig. 3D, box SD3). Because these genes were expressed in the absence of *Gtf2i* we concluded that these could be TFII-I (direct or indirect) target genes.

Genes upregulated in FO B cells in the absence of *Gtf2i* (105 cKO) were also expressed in other B cell subsets (Fig. 3E, box E1). For instance, a Pax5 target gene *Cecr2*, which is expressed during B cells development (*35*) is still expressed in FO B cells in the absence of *Gtf2i* (Fig. 3E). In contrast and unlike common and WT FO genes, the FO genes that are specific to the cKO group showed some enrichment of dendritic and macrophage subset genes such as *CD83* (Fig. 3E, box E2). Further, the genes that are expressed at low levels in the cKO FO subset were also expressed in other myeloid and stromal cell lineages (Fig. 3E, box E3). These observations indicate that TFII-I absence in FO B cells leads to expression of genes generally present during B cell developmental stages (*Vpreb3* and *Cecr2*) and genes which mark innate immune cells like *CD83* and *CD209a*.

#### MZ-specific genes

We next compared the transcriptomic features of MZ B cells in WT and cKO. The overlapping of 714 upregulated MZ specific genes in WT with 827 genes of *Gtf2i* cKO resulted in 626 common set of genes (Fig. 3F) that also exhibited comparable expression levels (Fig. 3G, left panel and Supp Table 3). These 626 MZ B cell genes are unaltered in the absence of TFII-I. When analyzed for gene features of other immune cells, 626 common MZ genes showed dominating MZ B cell features (Supp Fig. 3F, box SF1). These common genes also recapitulated features of innate immune cell genes (*33*) (Supp Fig. 3F, box SF2) and shared features characteristic of granulocytes and stromal cell compartments (Supp Fig. 3F, box SF3). While 201 MZ genes were upregulated in absence of *Gtf2i* (*Gtf2i* cKO MZ cell genes), including *Mpo, Gm4734, Mest, Gpr34* and *Asns* (Fig. 3G, right panel and Supp table 4). Likewise, 88 MZ genes were downregulated in *Gtf2i* lacking B cells (WT MZ cell genes) including *Zfp385a, Gap43*, and *E2f7*. WT MZ-specific genes (88 genes) didn’t demonstrate any significant features for other immune cell types (Supp Fig. 3G). The 201 genes derived from the MZ compartment of cKO demonstrated enrichment of genes belonging to developing stages of both B and T lymphocytes (Fig. 3H, box H1). These genes were generally poorly expressed in mature peripheral lymphocytes of WT mice (Fig. 3H, box H1, middle region dotted lines) and are mainly cell cycle related genes (Supp Table 4). Remarkably, in the absence of *Gtf2i*, MZ genes further exhibited features resembling innate immune compartments (Fig.3H, box H2). Additionally, cKO MZ genes showed commonality with granulocyte and stromal cell genes, which are generally not expressed in WT MZ B cells (Fig. 3H, box H3). Like FO B cells, these observations too suggest that there is an alteration of MZ defining basal transcriptome in the absence of TFII-I.

In summary, these transcriptomic data demonstrate that although in the absence of TFII-I the overall B cell FO and MZ differences are largely retained, B cells lacking TFII-I make FO B cells to exhibit signatures that skew them towards DC and macrophage subsets, while MZ B cells gained expression of cell cycle-related genes from developing stages of lymphocytes and further exhibit enhanced signatures characteristic of innate cells.

### Chromatin landscape of FO and MZ B cells in WT and *Gtf2i* cKO

Although TFII-I is a transcription factor that functions via cis-regulatory elements and expected to impact chromatin accessibility, this has not been directly explored. Open chromatin impressions are associated with cellular identities (*36*), thus to address whether absence of TFII-I would alter chromatin landscape of FO and MZ B cells, we performed Assay for Transposase-Accessible Chromatin with high-throughput sequencing (ATAC-seq) (*37*), using sorted FO and MZ B cells from WT and cKO spleens. After quality checks and alignment to the mouse genome, open chromatin regions (peaks) were called using MACS2 algorithm. We observed substantially fewer peaks in FO B cells in both WT and cKO (44,293 peaks in WT FO and 56,871 peaks in cKO FO) compared to MZ B cells (93,887 peaks in WT MZ and 104,543 peaks in cKO MZ) (Fig. 4A). Furthermore, we also noted that both FO and MZ B cell subsets of cKO have more open chromatin peaks than WT counterparts, (12,578 and 10,656) respectively. To determine where these peaks are located in the genome, we annotated the open chromatin peaks to five distinct genomic regions: promoter, exon, intron, TTS, and intergenic areas as shown in Fig. 4A. In the absence of TFII-I open promoter regions were reduced in cKO FO B cells while there was a gain of open chromatin peaks in the intergenic and intronic region. MZ B cell open chromatin location distribution in genome was similar between WT and cKO. We concluded that in the absence of TFII-I the chromatin is more permissive in both FO and MZ splenic B cells and that open chromatin of FO B cell in the absence of TFII-I is preferentially localized to the intergenic and intronic regions.

**Figure 4:**
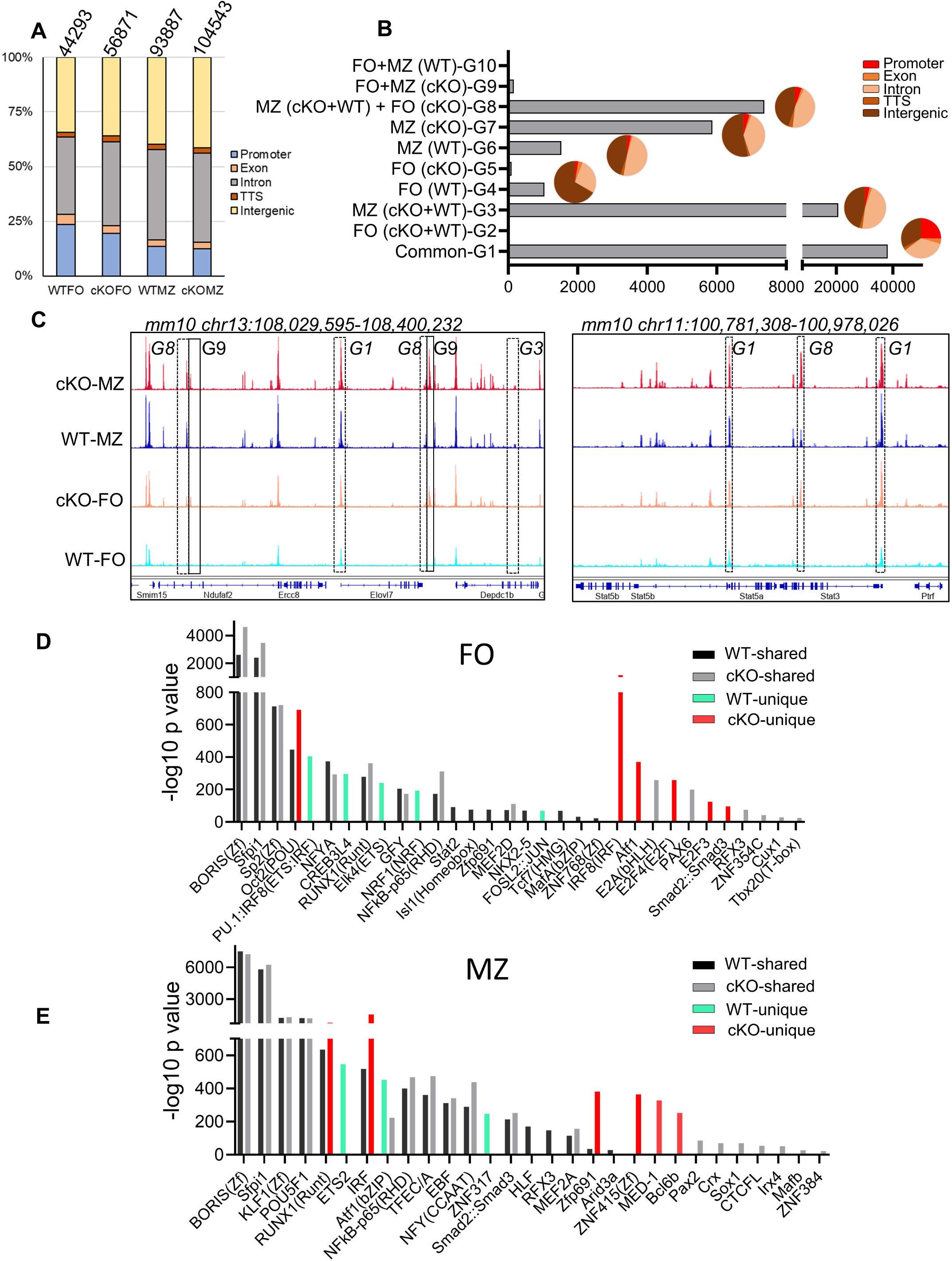
Chromatin landscape of FO and MZ B cells of WT and cKO mice. Quantitative description of chromatin landscape of four different splenic B cell populations: WT FO and MZ B cells and *Gtf2i* cKO FO and MZ B cells. (A) Peaks from FO and MZ B cells (numbers of peaks on top) from the indicated mouse strain and the annotations regarding their location in the genome. (B) Overlapping peaks from all four populations were identified using mergePeaks.pl suite (with -d 50) in HOMER. Ten different clustered groups (G1 – G10) of the overlapping peaks are shown with their indicated properties on the y axis. The pie charts show the peaks distribution in the genomic regions for the specified groups. (C) Genomic tracks of open chromatin (snapshot from IGV genome track viewer) from indicated B cell populations demonstrating open chromatin regions of chromosome13. Group specific peaks are outlined by boxes. (D - E) Presence of transcription factor motifs in open chromatin of WT and cKO FO (D) and MZ B cells (E) bars represent the -log10 p-value. cKO selective transcription factor motifs are marked by red color columns while WT selective transcription factor motifs are marked by light green color columns.

Open chromatin is often associated with presence or absence of specific transcription factor (TF) or families of TFs binding sites (*38*). To further understand the global effects of *Gtf2i* ablation on open chromatin landscape, we identified common and unique peaks using mergePeaks.pl from HOMER algorithm (*39*). We considered peaks across the cell types (FO or MZ) or strains (WT or cKO) as same if their start and end sites were within a span of 50 bp. This method provided 10 clustered groups with substantial number of peaks as defined in Fig. 4B. As expected, the largest number of peaks were common to both cell types (FO and MZ) and strains (WT and Gtf2i cKO) as defined in group G1. We hypothesized that these are the open chromatin features of mature splenic B cells. Second largest number of peaks was present in MZ B cells for both strains and is represented by group G3 (20694 peaks). These peaks represented distinct features of MZ cells compared to FO cells regardless of the strain. Furthermore, substantial number of unique peaks (5871) were present in cKO MZ (group G7), suggesting permissive chromatin of MZ B cells in the absence of TFII-I. Surprisingly, apart from these three major groups, 7370 unique peaks were also present in group G8 which is defined as a group consisting of peaks common to both FO and MZ of cKO and MZ of WT. We inferred that even FO B cells partly exhibit open chromatin features like that of MZ B cells in the absence of TFII-I.

Annotation of the common and unique peaks of these groups to the genomic regions is shown in the pie charts in Fig. 4B. The peaks in group G1 which represent mature B cell open chromatin are almost equally distributed between the promoter (25%), intronic (35%) and intergenic (33%) regions. However, the peaks in the MZ B cells specific group G3 fall mostly in the intronic and intergenic regions with less opening at the promoter region compared to G1. Like G3, the peaks of cKO MZ specific G7 and WT MZ specific G6 fall also mostly in the intronic and intergenic regions. Group G8 which denotes MZ like chromatin of FO B cells in the absence of TFII-I, shows open chromatin mostly in the intronic and intergenic regions, suggesting a role for intronic and intergenic region in regulating the chromatin landscape in these cells. Importantly, the changes in chromatin landscape in the absence of TFII-I become more evident when representative browser tracks of open chromatin peaks from chromosome 13 (Fig. 4C) are analyzed. Although these open chromatin features suggest that majority of chromatin landscape is not altered by ablation of TFII-I in both FO and MZ B cells, striking differences in selective peaks (as represented by G8) indicate that in the absence of TFII-I, a part of FO B cell open chromatin landscape features begin to skew towards chromatin features resembling that of MZ B cells.

To investigate the specific TF binding sites associated with ATAC-seq peaks, we analyzed the open chromatin of FO and MZ B cells identified in Fig. 4A using findMotifsGenome.pl algorithm of HOMER. Although many open chromatin features associated with WT and cKO enriched similarly in TF motifs in FO and MZ B cells (Fig. 4D, E), notably the IRF8(IRF) motif, which is mainly associated with myeloid and B lymphocyte differentiation (*40, 41*), was uniquely enriched in FO B cells of cKO (Fig. 4D). Furthermore, IRF and RUNX1 were enriched in both MZ of cKO and WT with higher enrichment in the cKO MZ than in the WT MZ (Fig. 4E). Presence of BORIS and Sfpi1 motifs were quite evident in both FO and MZ cell open chromatin from both strains. Similarly, the cKO MZ B peaks also additionally exhibit presence of Zfp691, ZNF415(Zf), MED-1 and Bcl6b motifs (Fig. 4E). We concluded that, while MZ B cell open chromatin expectedly shows presence of transcription factor motifs commonly found in innate immune cells, surprisingly in the absence of *Gtf2i*, FO B cells chromatin landscape tilts towards innate like features, notably exhibiting IRF motifs. These observations further confirm that part of the open chromatin of (both FO and MZ) B cells is altered in the absence of TFII-I, exhibiting additional enrichment of transcription factors binding motifs that are generally associated with innate cells such as IRF.

### FO and MZ defining open chromatin features in *Gtf2i* cKO B cells

After identifying basal state of open chromatin of FO and MZ B cell in the absence of TFII-I, we next analyzed FO and MZ defining open chromatin features in WT and cKO B cells, we used DiffBind algorithm (*42*). Significantly more chromatin peaks were open in MZ B cells compared to FO B cells of both strains, WT (39444 vs 983) and cKO (33222 vs 378) respectively (Fig. 5A). We further noticed >2fold decrease in differentially open chromatin (DOC) peak numbers in cKO FO B cells when compared to WT FO peaks (378 cKO FO vs 983 WT FO peaks) (Fig. 5A). When examined for their genomic location annotations, we found *Gtf2i* cKO FO and MZ specific peaks are less located at the promoter region compared to their WT counterpart while there was more cKO FO peaks at the intronic and intergenic region compared to WT FO peaks (Fig. 5B). These results indicate that TFII-I ablation also leads to reduced chromatin landscape differences between FO and MZ B cells.

**Figure 5:**
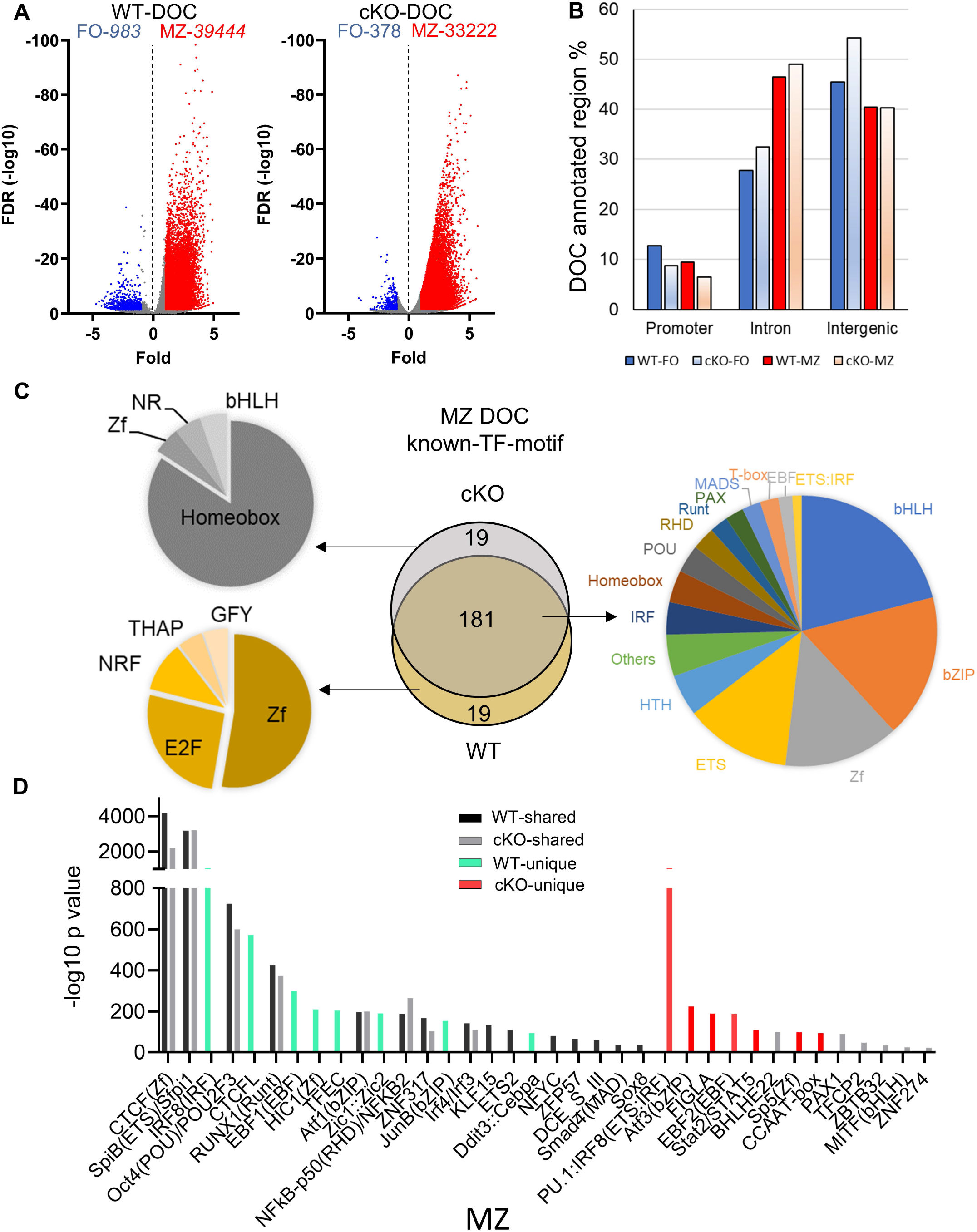
Transcription factors binding motifs in open chromatin of FO and MZ B cells. (A) Volcano plot showing the differentially open chromatin (DOC) from FO and MZ B cells of WT (left) and cKO (right); numbers of DOCs per cell type for the WT and cKO are shown on top. (B) The DOCs were annotated to the 3 genomic regions (promoter, Intron, Intergenic) and the percentages of these DOC annotated regions are shown for the four cell types. (C - D) The top 200 transcription factor (TF) motifs of DOC of MZ from WT and cKO B cells identified by known (C) and de novo (D) TF motif algorithm of HOMER. (C) Venn diagram (middle) shows the number of predicted common and unique TF binding motifs. Common motifs are represented by the pie chart on the right while the unique ones are represented on the left side. (D) bars represent the -log10 p-value. cKO selective transcription factor motifs are marked by red color columns while WT selective transcription factor motifs are marked by light green color columns.

Next, we focused on MZ specific regions for presence of transcription factor motifs using both known and de novo algorithms of HOMER suite (Fig. 5C, D). By comparing the top 200 transcription factor motifs (identified by known motifs algorithm) of MZ DOC between WT and cKO, 90% (181) of motifs were shared between both genotypes and were significantly enriched transcription factor motifs of basic-helix-loop-helix (bHLH), basic-leucine zipper (bZIP), zinc finger (Zf), and ETS family (Fig. 5C). In the absence of *Gtf2i*, 19 motifs under MZ DOC were dominated by Homeobox TF motifs while the WT MZ DOC were enriched for Zf and E2F motifs (Fig. 5C).

In our *de novo* TF motif algorithm analysis (which ranks the dominating TF motifs of the TF-family), we found that SpiB, CTCF, POU2F and RUNX1(Runt) motifs were present in MZ B cells open chromatin of both WT and cKO. Interestingly, IRF8, CTCFL, EBF1 and HIC1 motifs were selectively enriched in WT MZ specific open chromatin, while PU.1:IRF8, Aft3, EBF2 and Stat2/STAT5 motifs were dominated in the cKO MZ specific open chromatin. We noticed that some of the DOC features between MZ and FO, observed in WT B cells, such as IRF8 and HIC1 motifs in MZ DOCs were missing in cKO DOCs. Taking cues from previous analysis (Fig. 4), we speculated this may be due to pre-existing open chromatin in FO B cells of cKO and thus might not be reflective of differentially open chromatin analysis in cKO MZ B cells. These selective patterns of myeloid like TF-motifs enrichments suggest that the altered chromatin landscape may stem from FO B cell stages itself and the absence of TFII-I may influence all mature B cells.

### Innate response of B cells in absence of *Gtf2i*

It is well-established that murine B cells are less responsive to bacterial endotoxin LPS compared to the innate immune cells (e.g., macrophages and dendritic cells) (*43*). Given the clear indications from transcriptome and chromatin landscape analysis that removal of TFII-I accentuates innate-like features in B cells, we next aimed to test this notion by challenging B cells from WT and cKO *in vitro* with various dosages of LPS and assay for gene expression at various time points using Toll-like Receptor Signaling Pathway RT^2^ Profiler PCR Array (Fig. 6A). Naïve splenic B cells from WT and cKO mice were stimulated with 1 and 5 µg/ml of LPS for 0, 2, 8 and 24 hours. After RNA extraction and cDNA preparation, samples were subjected to the PCR array analysis. Differential gene expression analysis showed that 40% of the genes (34 genes) of the Toll-like receptor signaling pathway (84 genes) were expressed at higher levels in cKO B cells at basal level itself (Fig. 6B, red), while six out of the 84 genes exhibited higher expression in WT B cells (Fig. 6B, blue). These results show that at the basal level most of the Toll-like receptor pathway genes are upregulated in the cKO B cells compared to WT B cells.

**Figure 6:**
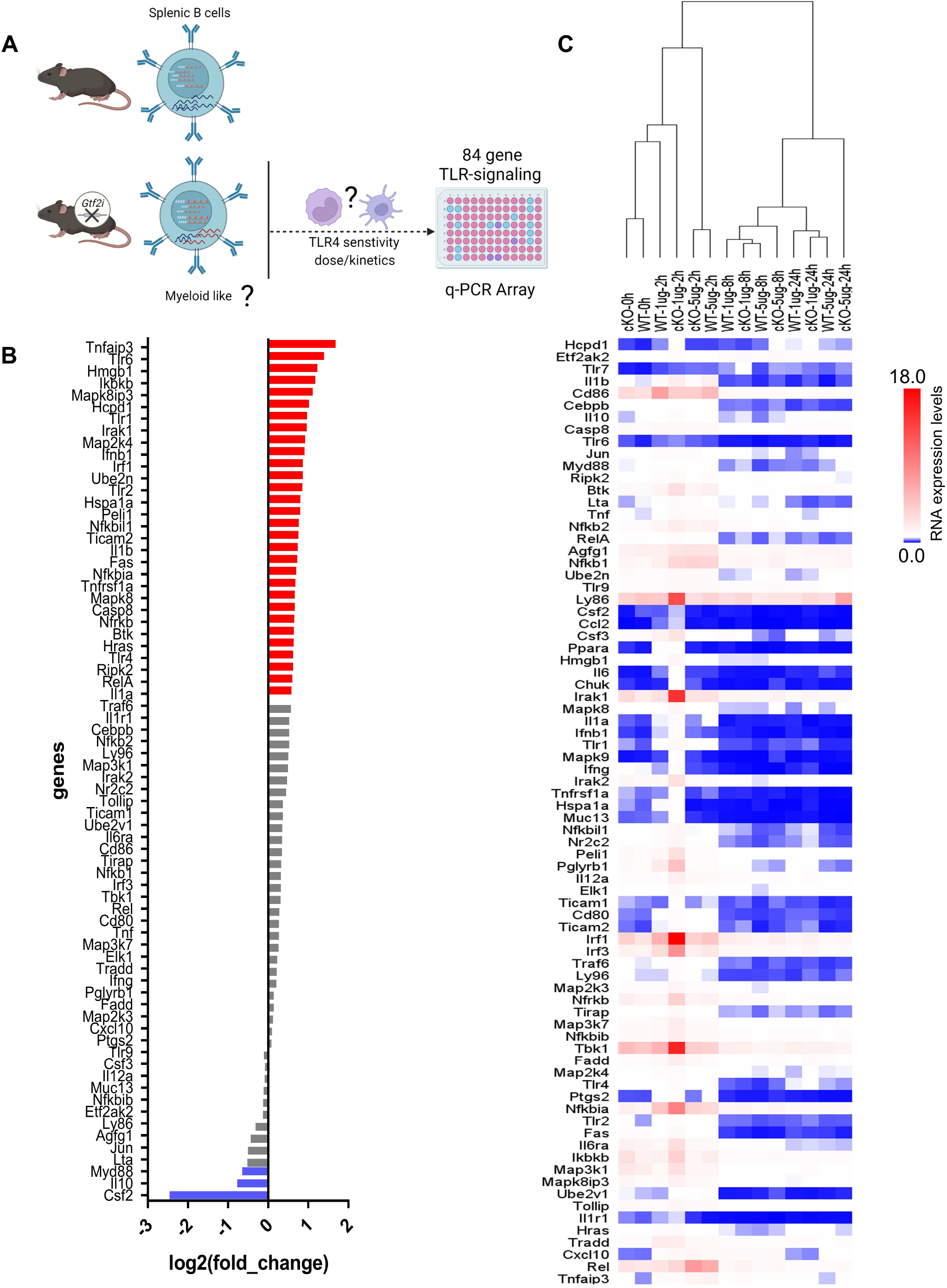
Transcriptome and chromatin landscape of LPS responsiveness splenic B cells upon Gtf2i ablation. Scheme representing the identified differences of transcriptome and open chromatin in B cells upon *Gtf2i* ablation. These differences indicate cKO B cells to possess myeloid like features, which are tested for TLR4 sensitivity by activation with LPS (5 and 1µg/ml) at early, mid and late time points (2, 8 and 24 hours) using TLR-signaling array. (B) Basal level expression of genes downstream of TLR-signaling in WT and cKO B cells. Red bars represent 30 genes highly expressed in cKO and blue are highly expressed in WT by at least 1.5-folds (n=4). (C) Heatmap representing the RNA expression levels of 84 TLR-responsive genes from LPS high (5ug/ml) and low (1ug/ml) dose at early (2h), mid (8h) and late (24h) time points (n=3). Relative expression of target genes was calculated to geometric mean of Ct values of *Gusb* and *Hsp90a*.

Interestingly, the suboptimal dose of LPS (1µg/ml) for B cells shows striking differences at early time point (2h). In hierarchal clustering the WT B cells clustered with unstimulated B cells while cKO B cells with the LPS 1µg/ml dose clustered away from WT and the unstimulated samples but closer to LPS 5µg/ml dose. We noticed with 1µg/ml dose activation at 2h in cKO B cells some genes, *Ly86, Irak1, Irf1, Tbk1* and *Nfkbia* were induced even higher than 5µg/ml dose. However, the response to optimal dose of LPS (5µg/ml) was similar between the cKO and WT B cells at earlier and late time points (2 and 24 hours) while mid time point (8h) shows some differences (Fig. 6C). Consistent with the RNA-seq and ATAC-seq analysis, these results together suggest that in the absence of TFII-I, the LPS sensitivity in B cells is increased, further indicating enhancement of their innate properties.

### Skewed B cell properties in absence of *Gtf2i*

Because our results showed that *Gtf2i* cKO MZ B cells have enhanced expression of genes involved in cell cycle regulation like that of developing lymphocytes (Fig. 3), we analyzed the proliferative properties of total splenic B cells after LPS stimulation. B cells from *Gtf2i* cKO demonstrated higher proliferation capacities at 72h after optimal 10µg/ml dose of LPS activation (Fig. 7A and quantified 7B). Similar differences were observed at suboptimal doses too (data not shown), while there was no difference when cells were stimulated through BCR crosslinking using anti-IgM Fab’2 (Fig. 7A). Remarkably, a far greater number of cells progressed into cell cycle in cKO B cells compared to WT B cells (Fig. 7B, G0 population). This increased proliferative property was further demonstrated by higher division index of cKO B cells at 72h (Fig. 7C).

**Figure 7:**
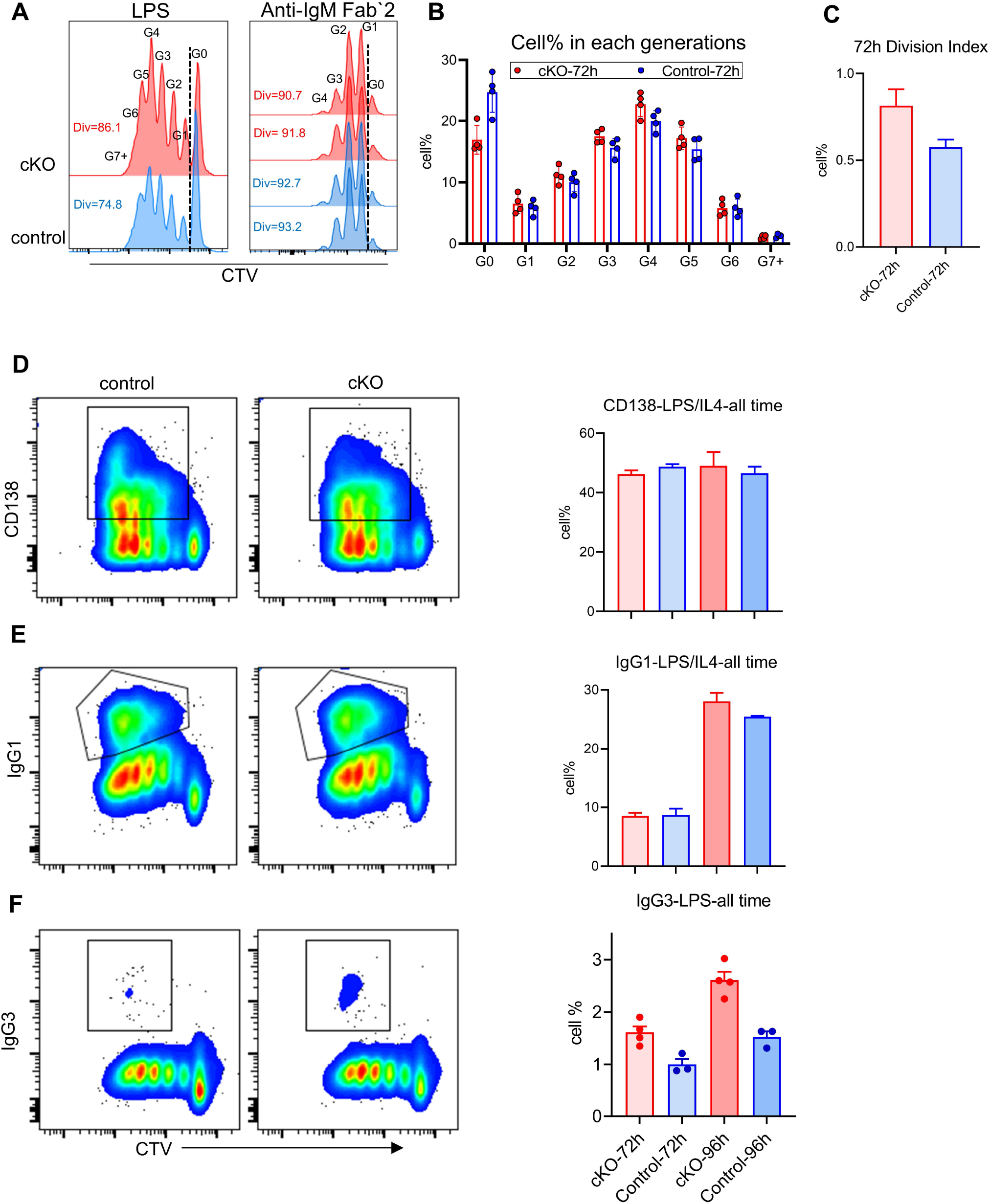
Functional characterization of splenic B cells from WT and cKo derived mice Splenic B cells from WT and cKO were activated with LPS (10ug/ml), F(ab’)2 anti-IgM (10ug/ml) or LPS (10ug/ml) plus IL-4 (20ng/ml) for the indicated time points followed by flow cytometry analysis. (A) Representative flow cytometry plot of Cell Trace Violet (CTV) dilution after 72 hours of LPS (n = 4) or anti-IgM (n = 2) activation. The Div numbers indicate the percentages of divided cells for the indicated mouse strain. Numbers above the CTV peaks refer to the cell generation (G0 – G6) (B) Cell percentages at each cell division after 72 hours of LPS activation. Bars represent the mean +/- SEM obtained from 4 independent experiments. (C) Division Index was calculated for cKO and WT B cells after 72 hours using FlowJo proliferation platform. (D - E) Representative flow cytometry plot of CD138 (D) or IgG1 (E) versus CTV for WT and cKO (left) after 96 hours of LPS plus IL-4 activation. Gating of CD138+ and IgG1+ cells are indicated. The bar plot on the right shows the average of the CD138+ or IgG1+ cell percentages at the indicated times. (F) Representative flow cytometry plot of IgG3 versus CTV after 96 hours of LPS activation and the bar plot shows the IgG3+ cell percentages at the indicated time points. Data represent 4 independent experiments.

To characterize these B cells functionally, LPS and IL4 treated splenic B cells from WT and cKO mice were tested for CD138^+^ plasma cells differentiation and IgG1 class switching. Although we did not observe any difference in plasma cell differentiation or IgG1 class switching (Fig. 7D-E), switching to inflammation associated IgG3 was significantly higher in cKO B cells (Fig. 7F). We concluded that in the absence of *Gtf2i*, splenic B cells become more responsive to innate signals like LPS activation that is reflected in enhanced proliferation and become biased towards class switching to a potent proinflammatory antibody, IgG3 isotype.

## Discussion

Immune responses to pathogens often involve a two-step process: first a nonspecific but rapid defense mechanism of the innate immune system is triggered and subsequently, a specific but slow defense mechanism of the adaptive immune system kicks in. While the innate arm is conserved from drosophila to man, the adaptive arm is exclusively present in vertebrates. These two arms evolved different surface receptors and specialized cells to respond distinctly to different cellular threats and environmental insults via triggering distinct intracellular signaling pathways (*44*). Although there is a boundary between these two arms with distinct cells and signaling pathways mediating such immune responses, murine MZ B cells are ideally located as sentinels at the interface between the circulation and lymphoid tissue to respond to blood-borne pathogens, thereby adopting a hybrid immune strategy that crosses over such boundaries (*19*). Thus, together with B-1 B cells, MZ B cells are endowed with a “natural memory” that provides a bridge between innate and adaptive immune responses (*20*). Consistent with this notion, MZ B cells are intrinsically capable of very rapidly maturing into plasmablasts, which may be due, in part, to the ability of MZ B cells to respond quickly than follicular B cells, to both B cell receptor (BCR) and to Toll-like receptors (TLRs) (*20*). In addition, MZ B cells also express high levels of TLRs (comparable to innate cells like DCs, macrophages and granulocytes). These properties allow the MZ B cells to crossover between the two arms of the immune system and rapidly respond to pathogens both via T-independent as well as T-dependent pathways (*17, 20*).

In this study, we determined that conditional ablation of *Gtf2i* in B cells leads to significant reduction in MZ B cells and a slight increase in FO B cells (Fig 1), without altering the number of splenocytes and B cells in spleen. Moreover, there was no appreciable difference in the viability of B cells with or without BAFF between control and *Gtf2i* cKO mice, perhaps suggesting a compensatory mechanism whereby reduction in MZ B cells is balanced by a concomitant increase in FO B cells. More surprising is the fact that the remaining MZ B cells as well as the FO B cells derived from *Gtf2i* cKO mice acquired accentuated molecular features of the innate-like properties of MZ B cells. This is reflected in total transcriptomic/RNA-seq (Figs 2 and 3) as well as chromatin landscape/ATAC-seq analyses (Figs 4 and 5). Although the exact molecular mechanisms for this function of TFII-I is currently unclear, there are several indications from our results. One key molecule may be the RNA binding protein (RBP) family gene. *Zfp36l2* was selectively upregulated in WT MZ B cells, but not in *Gtf2i* cKO MZ B cells (Fig 3A). ZFP36L1 protein post-transcriptionally represses expression of the transcription factors KLF2 and IRF8, which are known to favor the FO B cell phenotype. Given this established role of *Zfp36 family* in MZ B cell maintenance (*25*), downregulation of *Zfp36l2* might provide a rationale as to why the cKO mice have reduced numbers of MZ B and enhanced number of FO B cells. Consistent with this notion, we also noted that the IRF8 motif was uniquely enriched in FO B cells of cKO (Fig. 4). Given that TFII-I has been implicated in regulating homeotic genes like *Dlx5* and *Dlx6* (*45*) our observation that loss of TFII-I results in the open chromatin of homeobox containing motifs (Fig 5C) also suggests that regulation of homeobox genes in B cell function could be an important role of TFII-I. Further, it can also substantiate the association of homeobox transcription factors with MZ B cell lymphomagenesis (*46*) thereby indicating a potential role of TFII-I in this process.

B cell response to LPS is far less robust compared to innate cells, requiring greatly increased dose of LPS to elicit a comparable downstream effect (*43*). We rationalized that if *Gtf2i* cKO B cells acquired enhanced innate-like features, they might respond to a lower dose of LPS. We found that LPS sensitivity of these cells was also altered (Fig 6). In particular, a suboptimal dose of LPS at early time points elicited a robust response in cKO ex vivo splenic B cells, compared to the WT control B cells. To our surprise, 5 genes (*Ly86, Irak1, Irf1, Tbk1* and *Nfkbia*) were even more induced with the lower dose of LPS compared to higher dose. This apparently paradoxical regulatory mechanism may be due to time kinetics and/or dose dependent activation of stimulus specific downstream genes via enhancer-based mechanisms (*47*). Further specific studies are required to understand this interesting phenomenon.

To further distinguish functional differences between cKO and WT B cells and considering the fact the *Gtf2i* cKO MZ B cells exhibit enhanced expression of cell cycle genes like developing lymphocytes, we analyzed their proliferative potentials in response to LPS. B cells from *Gtf2i* cKO demonstrated higher proliferation capacities at 72h both at suboptimal and optimal dose of LPS (Fig 7A), while no such appreciable differences were noted when these cells were stimulated using anti-IgM Fab’2. Consistent with this observation, we also noted a far greater number of cells progressed into cell cycle in cKO B cells compare to WT B (Fig. 7B, G0 population) as demonstrated by higher division index of cKO B cells at 72h (Fig. 7C). Thus, a TFII-I deficiency in an adaptive immune cell such as B lymphocytes leads to a better proliferation capacity in response to innate signals. Differences in proliferative responses to TLR4 stimulation and BCR crosslinking also indicate the possibility that TFII-I may play critical role during T-independent B cell response rather than T-dependent one. Further, this gained cell number can potentially generate increased immune response, which could be beneficial or deleterious if not regulated properly. Paradoxically, TFII-I is noted to be an oncogene and a gain of function mutation is involved in thymomas and exhibits pro-proliferative capabilities in *in vitro* experiments (*12*), although proliferative potentials of TFII-I in B cells have never been tested before. Given the distinct and often opposing roles of TFII-I and its isoforms and its different family members in various cell type, further work needs to be done to clearly delineate the function of TFII-I in proliferation in different cell types.

Functional studies of CD138^+^ plasma cell differentiation and Ig class switching upon LPS and IL4 or only LPS stimulation *ex vivo* further confirm that in the absence of *Gtf2i*, splenic B cells exhibit enhanced responsiveness to LPS by undergoing increased proliferation (a characteristic feature of innate cells), without significantly altering their *in vitro* plasma differentiation properties. But interestingly, *Gtf2i* cKO B cells are clearly biased towards class switching to a potent proinflammatory antibody, IgG3 (*48*), further suggesting a role for TFII-I in altering B cell differentiation properties.

In these studies, we uncovered an unexpected cell-type specific role for TFII-I in murine B cell homeostasis through its conditional ablation. Given that the gain-of-function of TFII-I results in tumorigenic properties, we also conjecture that gain of TFII-I function could lead to B cell specific tumors. We further speculate that the B cell specific functions of TFII-I might be related to its known role in autoimmune disorders, like Sjogren syndrome and Lupus. Especially in light of the fact that serum IgG levels (including IgG3) are elevated in Sjogren syndrome and B cell subsets are altered (*49*), association of these features with lack of TFII-I could explain some aspects of the disease. Finally, although typical WBS syndrome clinical features do not include immune dysregulation, Kimura *et al* recently showed differential expression of genes related to B cell activation in WBS patients relative to controls (*50*). We anticipate future work will address the precise role of TFII-I in autoimmunity and WBS.

## Materials and Methods

### Mice

Gtf2i^fl/+^CD19-Cre+ (Het) and Gtf2i^fl/fl^CD19-Cre+ (HO) were generated by crossing Gtf2i^fl/fl^ (*21*) with Cd19-cre (006785, The Jackson Laboratory). BL6 (WT), Gtf2i^fl/+^CD19-Cre+ (Het) and Gtf2i^fl/fl^CD19-Cre+ (HO) mice were maintained in the animal facility of the National Institute on Aging. Eight to 12 weeks old mice were used for all experiments. The studies were carried out in accordance with the recommendations in the Guide for the Care and Use of Laboratory Animals (NRC 2010). Mice were euthanized with carbon dioxide and spleens were harvested for analysis. The protocol was approved by the Animal Care and Use Committee of the NIA Intramural Research Program, NIH. This program is fully accredited by the Association for Assessment and Accreditation of Laboratory Animal Care International (AAALAC) (File 000401), registered by the United States Department of Agriculture (51-F-0016) and maintains an assurance with the Public Health Service (A4149-01).

### MZ, FO and B cell isolation

Primary B lymphocytes were isolated from spleens of WT, Het and HO mice using EasySep B cell kits by negative selection according to the manufacturer’s instructions (Stemcell Technologies, Canada). B cells were stained with anti CD21-FITC and anti CD23-PE antibodies (Biolegend, San Diego, CA) and sorted at the BD FACS-Aria Fusion. Follicular (FO) B cells were sorted as CD21^lo^ CD23^hi^ and Marginal Zone (MZ) B cells were sorted as CD21^hi^ CD23^lo^. Cells were cultured in RPMI 1640 medium (Invitrogen, MA) containing 10% heat inactivated FBS (Gemini Bioproducts, CA), 50μM β-mercaptoethanol (Sigma-Aldrich, MO), 1% L-glutamine and 1% penicillin-streptomycin solution (Gibco, MA). B cell purity was > 95% based on flow cytometric analysis following staining with anti-CD19 (Biolegend, San Diego, CA). FO and MZ B cells were >90% pure based on the criteria that they were sorted on.

### Transcriptomic analysis (RNA-Seq)

Total RNA was extracted from FO and MZ B cells using the RNeasy Mini Kit (Qiagen, Valencia, CA) according to the manufacturer’s protocol. Total RNA was sequenced at the Johns Hopkins Deep Sequencing and Microarray Core using standard protocol for NextSeq 500 sequencer. Ribosomal RNA was depleted, barcoded libraries were made, and 50bp single end reads were generated for each sample. Reads were analyzed using Galaxy RNA-seq pipeline (usegalaxy.org) and adapter trimmed sequences were aligned to mouse genome (mm10) using HISAT2. FeatureCounts and htseq-count were used for differential gene expression analysis using DESeq2.

### ATAC-seq

FO and MZ splenic B cells were FACS sorted and ATAC-seq libraries were made as previously described (Buenrostro, Giresi et al. 2013) with minor modifications. Briefly, 100,000 cells were lysed in 100 ul of lysis buffer (10 mM Tris-Cl pH 7.4, 10 mM NaCl, 3 mM MgCl_2_, 0.1% NP-40). After centrifuging at 500g for 10 min at 4°C, pelleted nuclei were resuspended with 50 ul of transposition mix (1X Tagment DNA buffer, Tn5 Transposase, nuclease-free H_2_O) and incubated for 30 min at 37°C in a thermomixer with slow revolutions. Transposed DNA was purified using MinElute columns (28004, QIAGEN) and subsequently amplified with Nextera sequencing primers and NEB high fidelity 2X PCR master mix (New England Biolabs) for 11 cycles. PCR-amplified DNA libraries were size selected after fractioning on high melting agarose gel electrophoresis. Library was pooled and sequenced using the Illumina NextSeq sequencer with paired end reads of 75 bases. After sequencing the ATAC-seq analysis was performed on Galaxy server (usegalaxy.org). Reads were run for quality control and treated accordingly for trimming and adapter removing. QC passed reads, were aligned to mouse mm10 genome using Bowtie2 and peak calling was doing using MACS2 peak caller. Differential open chromatin was identified using DiffBind algorithm.

### Activation and Gene expression analysis

Splenic B cells were isolated from WT and cKO mice and cultured in presence of LPS (InvivoGen, San Diego, CA) 5 µg/ml or 1 µg/ml for 2, 8 and 24 h time points. Total RNA was extracted from B cells using RNeasy Mini Kit (Qiagen, Valencia, CA). 100ng RNA was reverse transcribed using SuperScript IV VILO Master Mix (Thermofisher Scientific, MA) and the cDNA was loaded onto the “Mouse Toll-Like Receptor Signaling Pathway” RT^2^ Profiler PCR Array according to the manufacturer instructions (Qiagen, Valencia, CA). Expression of the target genes were normalized to the geometric mean of Ct values of the two housekeeping genes *Gusb* and *Hsp90a* using the ddCt method (n = 3). Basal level (0 h) fold change was calculated by dividing the gene expression of cKO B cells on the gene expression of WT B cells (n = 4).

### Proliferation and class switching

Splenic B cells were labeled with Cell Trace Violet (CTV) (Invitrogen, MA) according to the manufacturer instructions. After washing cells were activated with LPS 10 µg/ml (n = 4) or with F(ab’)2 anti-IgM 10 µg/ml (n = 2) for 72 h followed by flow cytometry analysis for CTV dilution. Each division generation are labeled from G0 – G6. Proliferation plugin of FlowJo (TreeStar BD Biosciences) was used to calculate division and expansion index. For plasma cell differentiation and IgG1 class switching analysis, LPS/IL4 activated B cells were co-stained with anti-CD138-BV711 and anti-IgG1-PE respectively at 72 and 96 h time points. For IgG3 class switching analysis, LPS activated B cells were stained with anti-IgG3-FITC at 72 and 96 h. Flow acquisition was performed on BD Symphony A5 system and analysis was done on FloJo (TreeStar BD Bioscience).

