## Supplementary Figures for "Transcription factor TFII-I fine tunes innate properties of B lymphocytes"

**Supp. Fig1:**

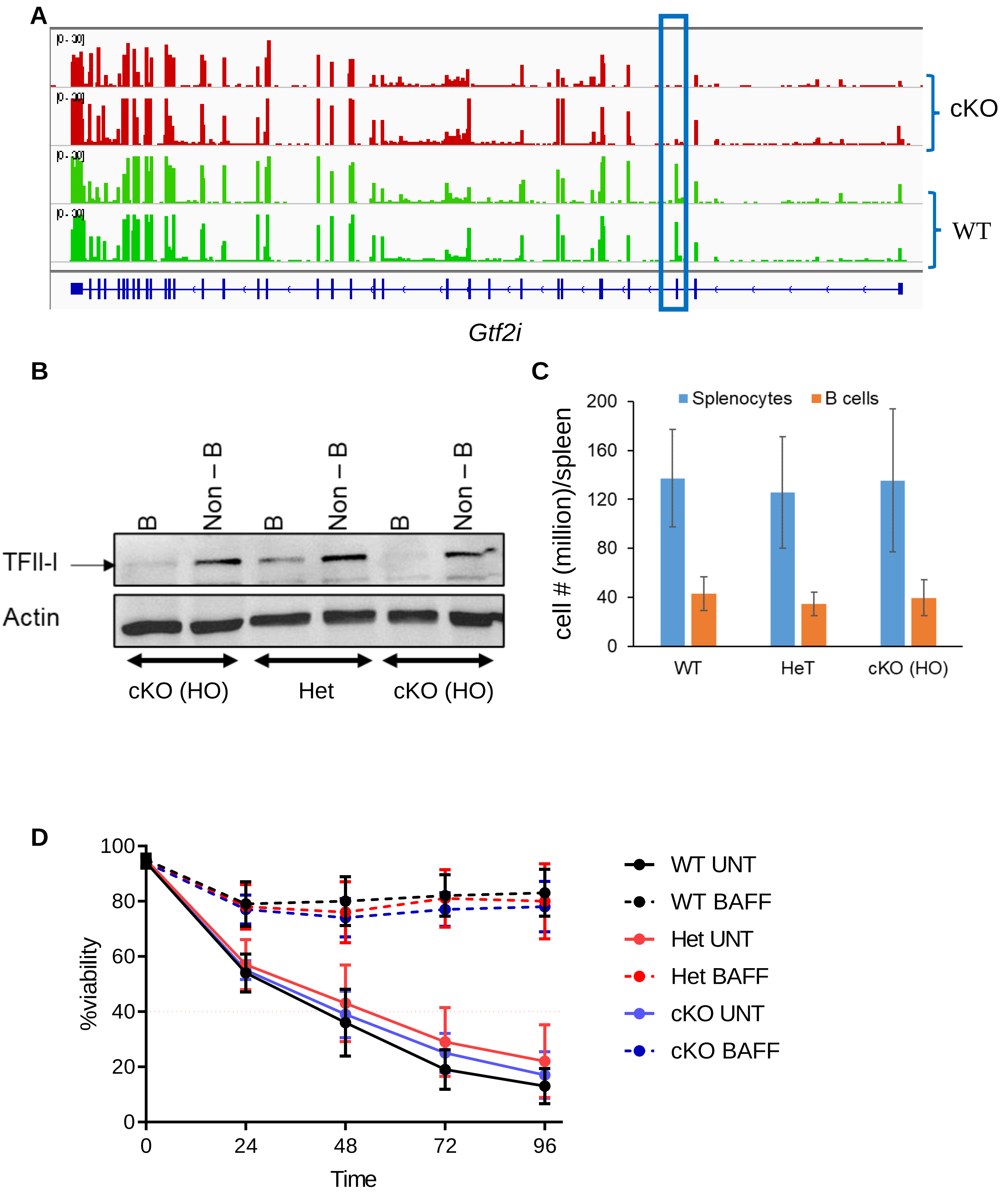
A
cKO
WT
Gtf2i
B
B
Non – B
B
Non – B
B
Non – B
TFII-I
Actin
cKO (HO)
Het
cKO (HO)
C
Splenocytes
B cells
cell # (million)/spleen
200
160
120
80
40
0
WT
HeT
cKO (HO)
D
%viability
100
80
60
40
20
0
0
24
48
72
96
Time
WT UNT
WT BAFF
Het UNT
Het BAFF
cKO UNT
cKO BAFF

**Supp. Fig1:**

**A:** Genome Browser tracks (IGV) from the RNA-Seq analysis of *Gtf2i gene in* WT and cKO B cells. Figure shows duplicate samples of each strain. Box represent exon 3 position of the Gtf2i gene.

**B:** Splenic B cells were isolated from the *Gtf2i*$^{fl/fl}$*CD19-Cre+ cKO (*HO) and *Gtf2i*$^{fl/+}$*CD19-Cre* Het mice using Easysep B cell isolation kit by negative selection and the rest of the splenocytes were used as control (Non-B cells). B and non-B cells were lysed in RIPA lysis buffer. Whole cell extracts were fractionated by SDS-PAGE and TFII-I protein expression was analyzed by immunoblotting. β-actin was used to normalize between samples.

**C:** Total splenocytes and splenic B cell numbers from cKO (HO), Het and WT mice. Data are representative of 6 independent experiments.

**D:** Splenic B cells from cKO (HO), Het and WT mice were cultured ex vivo at 37 °C with or without BAFF (200ng/ml) for the indicated times. Viability was determined by propidium iodide staining and flow cytometry. Error bars represent the standard error of the mean between experiments and data are representative of three independent experiments.

**Supp. Fig. 2**

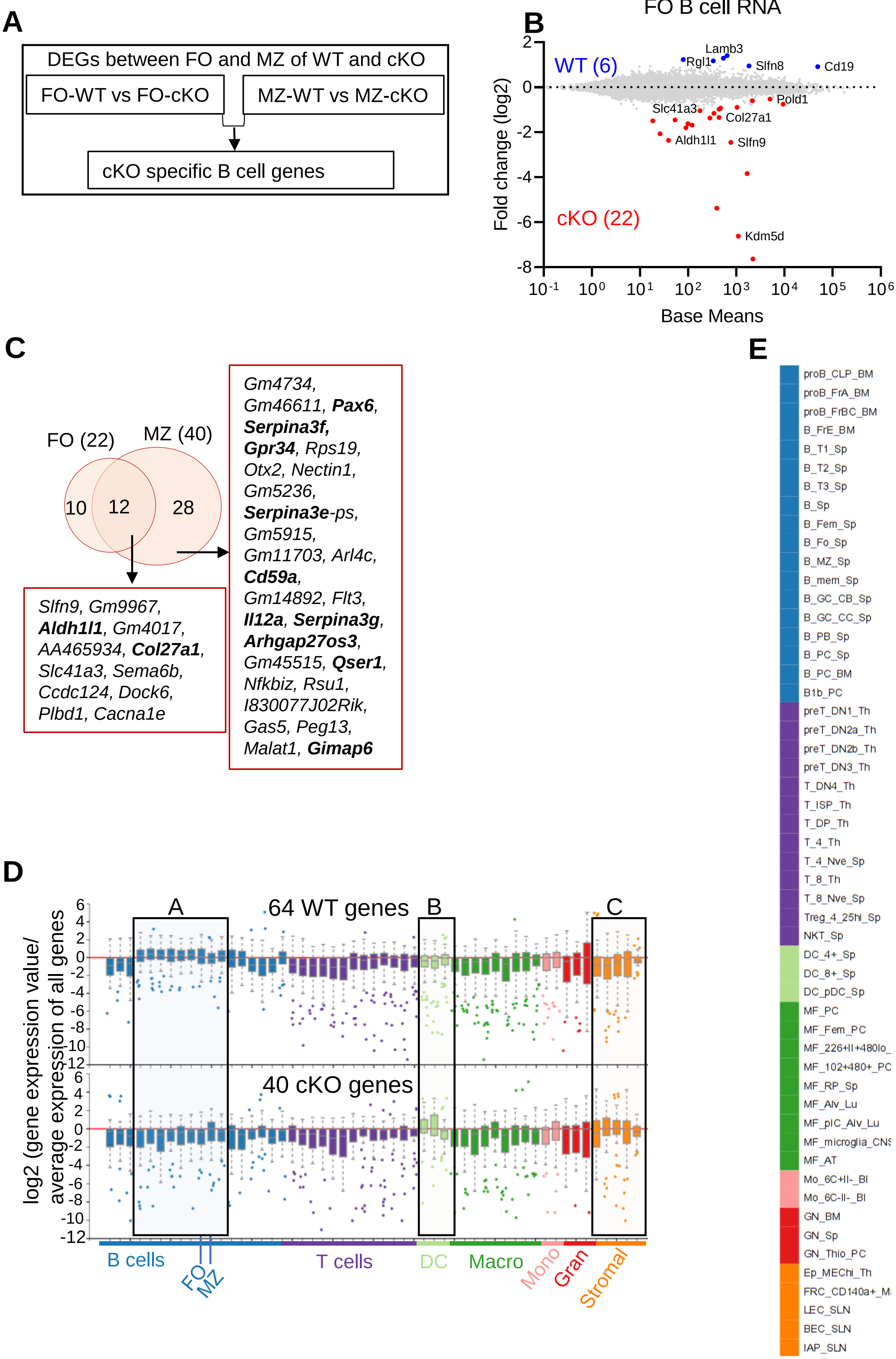
A
DEGs between FO and MZ of WT and cKO
FO-WT vs FO-cKO
MZ-WT vs MZ-cKO
cKO specific B cell genes
B
FO B cell RNA
WT (6)
cKO (22)
Lamb3
Rgl1
Slfn8
Cd19
Pold1
Slc41a3
Col27a1
Aldh1l1
Slfn9
Kdm5d
Fold change (log2)
Base Means
C
FO (22)
MZ (40)
10
12
28
Slfn9, Gm9967, Aldh1l1, Gm4017, AA465934, Col27a1, Slc41a3, Sema6b, Ccdc124, Dock6, Plbd1, Cacna1e
Gm4734, Gm46611, Pax6, Serpina3f, Gpr34, Rps19, Otx2, Nectin1, Gm5236, Serpina3e-ps, Gm5915, Gm11703, Arl4c, Cd59a, Gm14892, Flt3, Il12a, Serpina3g, Arhgap27os3, Gm45515, Qser1, Nfkbiz, Rsu1, I830077J02Rik, Gas5, Peg13, Malat1, Gimap6
D
64 WT genes
40 cKO genes
A
B
C
log2 (gene expression value/ average expression of all genes
B cells
FO
MZ
T cells
DC
Macro
Mono
Gran
Stromal
E
proB_CLP_BM
proB_FrA_BM
proB_FrBC_BM
B_FrE_BM
B_T1_Sp
B_T2_Sp
B_T3_Sp
B_Sp
B_Fem_Sp
B_Fo_Sp
B_MZ_Sp
B_mem_Sp
B_GC_CB_Sp
B_GC_CC_Sp
B_PB_Sp
B_PC_Sp
B_PC_BM
B1b_PC
preT_DN1_Th
preT_DN2a_Th
preT_DN2b_Th
preT_DN3_Th
T_DN4_Th
T_ISP_Th
T_DP_Th
T_4_Th
T_4_Nve_Sp
T_8_Th
T_8_Nve_Sp
Treg_4_25hi_Sp
NKT_Sp
DC_4+_Sp
DC_8+_Sp
DC_pDC_Sp
MF_PC
MF_Fem_PC
MF_226+II+480lo_
MF_102+480+_PC
MF_RP_Sp
MF_Alv_Lu
MF_pIC_Alv_Lu
MF_microglia_CNS
MF_AT
Mo_6C+II-_Bl
Mo_6C-II-_Bl
GN_BM
GN_Sp
GN_Thio_PC
Ep_MEChi_Th
FRC_CD140a+_M
LEC_SLN
BEC_SLN
IAP_SLN

**Supp. Fig. 2:**

A- Scheme demonstrating the comparison profile for the identification of differentially expressed genes (DEGs). FO genes in the WT were compared to the FO genes in the cKO and  WT MZ genes were compared to the cKO MZ genes.

B- MA plot displaying the log fold-change compared with mean expression and showing the DEGs in FO B cells of the WT (6 genes) and of the cKO mice (22 genes).

C- Venn diagram showing the overlapping of the  40 cKO MZ genes with the 22 cKO FO genes results in 12 genes that are *Gtf2i* cKO B cells specific. A list of the 12 *Gtf2i* cKO unique genes and the 28 cKO MZ unique genes are shown.

D- Box and whiskers plot shows the WT 64 MZ genes and the cKO 40 MZ genes and their expression levels in all the immune cells using ImmGen-database. Genes that are enriched in B cell compartments are shown in box A, genes that are enriched in DC are shown in box B while the genes that are enriched in granulocytes and stromal cells are shown in box C.

E- Scheme showing the cell populations as presented by ImmGen-Database and shown in all the heatmaps that are created from ImmGen-Database in 3B, 3C and Supp. Fig 4C.

**Supp. Fig. 3:**

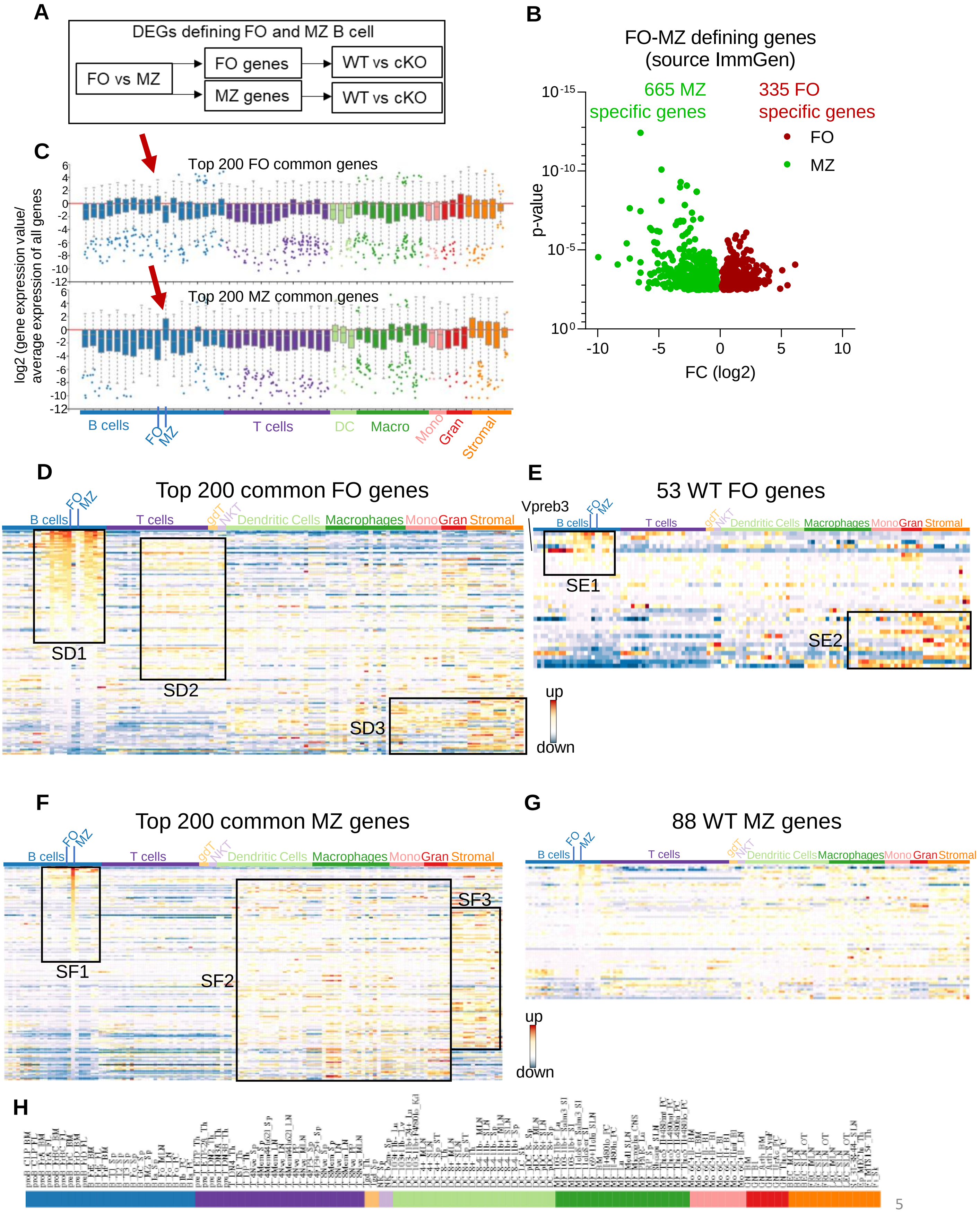
A
DEGs defining FO and MZ B cell
FO vs MZ
FO genes
MZ genes
WT vs cKO
WT vs cKO
B
FO-MZ defining genes
(source ImmGen)
665 MZ
specific genes
335 FO
specific genes
FO
MZ
p-value
FC (log2)
C
Top 200 FO common genes
Top 200 MZ common genes
log2 (gene expression value/
average expression of all genes
B cells
FO
MZ
T cells
DC
Macro
Mono
Gran
Stromal
D
Top 200 common FO genes
SD1
SD2
SD3
E
53 WT FO genes
Vpreb3
SE1
SE2
up
down
F
Top 200 common MZ genes
SF1
SF2
SF3
G
88 WT MZ genes
Dendritic Cells
Macrophages
gdT
NKT
H

**Supp. Fig. 3:**

A: Scheme demonstrating the comparison profile for the identification of differentially expressed genes (DEGs) between FO and MZ genes in the WT and cKO mice. First FO genes were compared to the MZ genes per strain (WT or cKO). Second the WT FO genes were compared to the cKO FO genes, similarly WT MZ genes were compared to the cKO MZ genes.

B: Volcano plot showing the FO vs MZ genes of C57BL/6 (WT) mice from the Immunological Genome Database (ImmGen-Database). Data shows 335 FO specific genes versus 665 MZ specific genes.

C - D: Box and whiskers plot (C) and heatmap (D) shows the top 200 common FO genes (common in WT and cKO) and their expression levels in all the immune cells using ImmGen-database. Genes that are enriched in B cell compartments are shown in box SD1, genes that are enriched in T cell compartments are shown in box SD2 while the genes that are enriched in monocytes, granulocytes and stromal cells are shown in box SD3.

E: Heatmap showing the expression levels of the 53 WT specific FO genes in the different immune cells when they were input in the ImmGen-Database. Box SE1 shows the expression levels of these genes in the different subsets of B cells. Box SE2 shows their expression levels in the monocytes, granulocytes and stromal cells.

F: Heatmap like in D but for the top 200 common MZ genes. SF1 box shows the enrichment of these genes in the B cell compartment, SF2 shows that these genes are enriched in NK cells, dendritic cells, macrophages and monocytes. SF3 shows the subset of genes that are enriched in the granulocytes and stromal cells.

G: Heatmap showing the expression levels of the 88 WT MZ specific genes when they were input in ImmGen-Database.

H: scheme showing the cell populations as presented by ImmGen-Database and shown in all the heatmaps that are created from ImmGen-Database (Fig. 2E, 2H, Supp Fig. 2D – 2G).
